# Sulfide oxidation promotes hypoxic angiogenesis and neovascularization

**DOI:** 10.1101/2023.03.14.532677

**Authors:** Roshan Kumar, Victor Vitvitsky, Proud Seth, Harrison L. Hiraki, Hannah Bell, Anthony Andren, Rashi Singhal, Brendon M. Baker, Costas A. Lyssiotis, Yatrik M. Shah, Ruma Banerjee

## Abstract

Angiogenic programming in the vascular endothelium is a tightly regulated process to maintain tissue homeostasis and is activated in tissue injury and the tumor microenvironment. The metabolic basis of how gas signaling molecules regulate angiogenesis is elusive. Herein, we report that hypoxic upregulation of NO synthesis in endothelial cells reprograms the transsulfuration pathway and increases H_2_S biogenesis. Furthermore, H_2_S oxidation by mitochondrial sulfide quinone oxidoreductase (SQOR) rather than downstream persulfides, synergizes with hypoxia to induce a reductive shift, limiting endothelial cell proliferation that is attenuated by dissipation of the mitochondrial NADH pool. Tumor xenografts in whole-body WB^Cre^SQOR^fl/fl^ knockout mice exhibit lower mass and reduced angiogenesis compared to SQOR^fl/fl^ controls. WB^Cre^SQOR^fl/fl^ mice also exhibit reduced muscle angiogenesis following femoral artery ligation, compared to controls. Collectively, our data reveal the molecular intersections between H_2_S, O_2_ and NO metabolism and identify SQOR inhibition as a metabolic vulnerability for endothelial cell proliferation and neovascularization.

**Highlights:**

- Hypoxic induction of •NO in endothelial cells inhibits CBS and switches CTH reaction specificity
- Hypoxic interruption of the canonical transsulfuration pathway promotes H_2_S synthesis
- Synergizing with hypoxia, SQOR deficiency induces a reductive shift in the ETC and restricts proliferation
- SQOR KO mice exhibit lower neovascularization in tumor xenograft and hind limb ischemia models

## Introduction

Angiogenesis is a complex process that is essential for embryonic development and for wound repair in adults^1^. Dysregulation of this process is associated with the pathogenesis of infectious, malignant, ischemic, inflammatory and immune diseases^2,3^. Despite the anticipated promise of early antiangiogenic therapeutics for treating disorders ranging from cancer to blindness^4^, these drugs have shown limited efficacy, highlighting the need for alternative and preferably, combination therapies to counteract issues with resistance and/or neovascularization via other pathways^5^. Hydrogen sulfide (H_2_S) is a pro-angiogenic metabolite^6^, which interacts with the nitric oxide (•NO)-dependent cGMP signaling pathway^7^, and increases endothelial cell growth and migration via phosphorylation of ERK, Akt and p38^8^. Vascular endothelial growth factor (VEGF) reportedly stimulates H_2_S release^8^. On the other hand, sulfur amino acid restriction also serves as a pro-angiogenic trigger, paradoxically increasing H_2_S synthesis in addition to VEGF expression, signaling via the GCN2/ATF4 amino acid starvation pathway^9^. The mechanisms underlying •NO-dependent regulation of H_2_S biogenesis in endothelial cells and the role of sulfide oxidation in angiogenesis are however, poorly understood.

H_2_S is a product of the transsulfuration pathway enzymes, cystathionine β-synthase (CBS) and γ-cystathionine (CTH)^10^ (Figure 1A). The transsulfuration pathway plays a key role in regulating levels of the thrombogenic amino acid homocysteine, as well as in furnishing cysteine, the limiting substrate for glutathione synthesis^11,12^. The pathway is also a source of H_2_S, which inhibits complex IV and modulates cellular energetics^13,14^. Canonical roles of CBS and CTH lead to sulfur transfer from homocysteine to serine, forming cystathionine, which is subsequently metabolized to cysteine, a-ketobutyrate and ammonia. Addtionally, CBS and CTH exhibit substrate promiscuity, leading to noncanonical reactions that generate H_2_S from cysteine and/or homocysteine^15^. The regulatory heme cofactor in CBS sensitizes it to inhibition by carbon monoxide (CO) or •NO, decreasing cystathionine synthesis^16,17^, which in turn, promotes H_2_S synthesis by CTH^18^. Therefore, conditions that stimulate CO or •NO production can stimulate H_2_S synthesis^18^.

**Figure 1.**
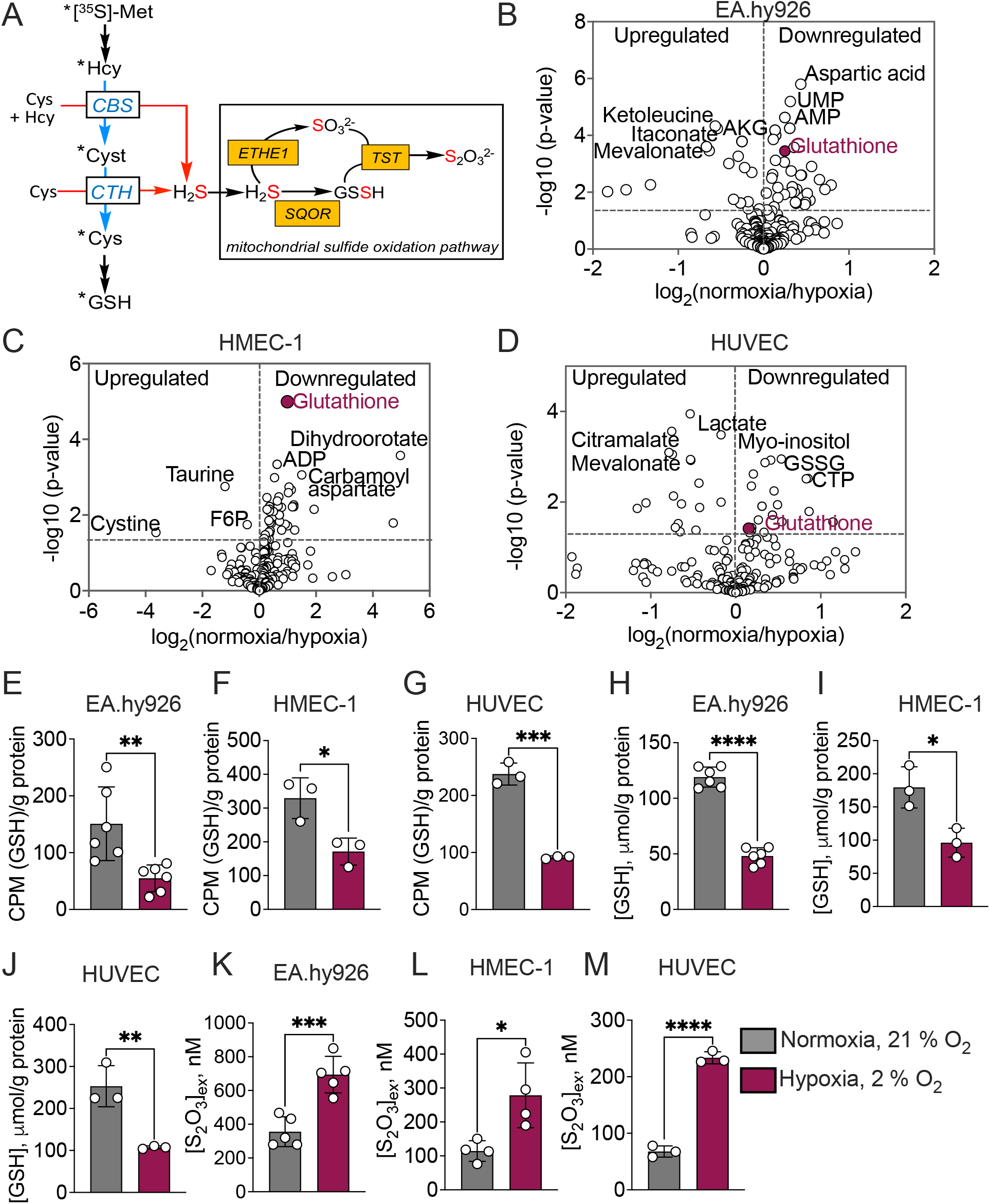
Hypoxic track switching in the transsulfuration pathway. (**A**) Scheme showing radiolabel transfer from [^35^S]-methionine to GSH (asterisks) via the canonical transsulfuration pathway (*blue arrows*), H_2_S synthesis (*red arrows*) and the mitochondrial oxidation reactions. Cyst, TST and ETHE1 are cystathionine, thiosulfate sulfurtransferase and persulfide dioxygenase, respectively. (**B-D**) Volcano plots reveal that GSH is significantly decreased in EA.hy926 (B), HMEC-1 (C), and HUVEC (D) cells grown under hypoxia versus normoxia (n=4). (**E-G**) [^35^S]-methionine incorporation into GSH is lower in hypoxia versus normoxia in the indicated cell lines. n = 3-6; *p=0.02, **p=0.007, and ***p=0.0002. (**H-J**) GSH levels are lower in the indicated cell lines. n = 3-6; *p=0.02 and **p=0.007, and ****p<0.0001. **(K-M**) Extracellular thiosulfate levels are higher in the indicated cell lines under hypoxia versus normoxia. n =3-5. *p= 0.017, ***p=0.0006, and ****p<0.0001. Samples in E-M were collected either at 24 h (Ea.hy926, HMEC-1) or at 16 h (HUVEC). Data represent mean ± S.D.

Low steady-state tissue H_2_S concentrations^19^ result primarily from the high activity of the mitochondrial sulfide oxidation pathway^20^. Sulfide quinone oxidoreductase (SQOR) commits H_2_S to oxidation (Figure 1A) while concomitantly reducing coenzyme Q (CoQ), which enters the electron transport chain (ETC) at the level of complex III. In addition to shielding cells from respiratory poisoning^21^, SQOR modulates electron flux in the ETC, causing a reductive shift in the mitochondrial NAD^+^ and CoQ pools^22^, activating aerobic glycolysis^23^ and glutamine-dependent lipid synthesis^24^. Interestingly, the pro-angiogenic effect of H_2_S is reportedly mediated in part by ETC inhibition and enhanced aerobic glycolysis^9^. While endothelial cells can meet up to 85% of their ATP needs from glycolysis^25^, complex III ablation decreases proliferation and angiogenesis and is associated with decreased amino acid levels^26^.

At H_2_S concentrations that inhibit forward electron transfer, cells prioritize its clearance by rerouting electrons through complex II using fumarate as a terminal electron acceptor^27^. A similar remodeling of the ETC via fumarate-dependent complex II reversal is observed in hypoxia^28^. Sulfide oxidation by SQOR protects against hypoxia-induced injury in brain, which is particularly vulnerable to O_2_ deprivation^29^. SQOR inhibition reportedly protects against heart failure in a mouse model^30^.

In the present study, we demonstrate that hypoxic upregulation of •NO synthesis in endothelial cells leads to metabolic reprogramming of the transsulfuration pathway via allosteric inhibition of cystathionine β-synthase, which shifts the reaction specificity of γ-cystathionase to H_2_S biogenesis. The proangiogenic effects of H_2_S depend on its oxidative metabolism as SQOR deficiency decreases cell proliferation, sprouting capacity, and tube formation in hypoxic endothellial cells. Whole-body SQOR knockout (KO) diminished tumor growth and angiogenesis in a xenograft model. Consistent with this data, SQOR KO mice demonstrated decreased muscle angiogenesis in a hind limb ischemia model induced via femoral artery ligation. This study identifies SQOR as a therapeutic target for inhibiting the pro-angiogenic effects of H_2_S under hypoxia.

## Results

### Hypoxia stimulates H_2_S biogenesis

Metabolomics analysis revealed decreased glutathione (GSH) in transformed (Ea.hy926 and HMEC-1) and primary (HUVEC) endothelial cells cultured under hypoxic (2% O_2_) versus normoxic (21%) conditions (Figure 1B-D, Table S1). To test our hypothesis that hypoxia redirects cysteine from GSH towards H_2_S synthesis, radiolabel transfer from [^35^S]-methionine to GSH was monitored (Figure 1A). Hypoxia decreased the magnitude of radiolabel incorporation as well as the GSH pool size in all three lines (Figure 1E-J). In contrast, extracellular thiosulfate, a stable biomarker of H_2_S metabolism, increased under hypoxia (Figure 1K-M). Three of four non-endothelial cell lines tested (HEK293, 143B and HepG2) also exhibited decreased radiolabel transfer and lower GSH pool size (Figure S1A-C and E-G), while significant differences were not observed in SH-SY5Y cells (Figure S1D,H). Hypoxic thiosulfate accumulation was only seen in HepG2 cells (Figure S1), indicating that decreased flux to GSH is not necessarily coupled to increased H_2_S metabolism under hypoxia. The complex vascular cell basal medium used for HUVEC culture, suppressed thiosulfate accumulation and GSH labeling although a diminution in the GSH pool was still observed (Figure S2A-C). The vascular cell basal medium also suppressed hypoxic thiosulfate accumulation by EA.hy926, suggesting that growth factors influence H_2_S homeostasis (Figure S2D). Regulation of CBS by nutrients and basic fibroblast growth factor has been reported previously^31^.

### •NO regulates hypoxic H_2_S homeostasis

Hypoxia increases eNOS activity and •NO production^32–35^. We therefore posited that •NO would indirectly induce H_2_S synthesis by CTH (Figure 2A). DETA NONOate, an •NO donor, elicited a dose-dependent increase in extracellular thiosulfate, while cPTIO, an •NO scavenger, decreased thiosulfate accumulation under normoxia (Figure 2B,C). Cystathionine, used to competitively inhibit H_2_S synthesis by CTH, also decreased hypoxic thiosulfate accumulation (Figure 2D), while arginine, a substrate for •NO synthesis, had no effect (Figure S3A). Cystine and homocystine, which upon intracellular reduction to cysteine and homocysteine, respectively, are substrates for H_2_S synthesis, increased extracellular thiosulfate under normoxia (Figure 2E,F). EA.hy926 (scrambled control) cells consumed extracelluar cystine more rapidly at 8h under hypoxia versus normoxia (Figure 2G). In contrast, partial knockdown of eNOS or the upstream regulators, HIF1α and HIF2α (Figure S3B,C) decreased hypoxic thiosulfate accumulation (Figure 2H,I) while roxadustat (FG4592), which stabilizes HIF, increased thiosulfate accumulation under normoxia (Figure S3D).

**Figure 2.**
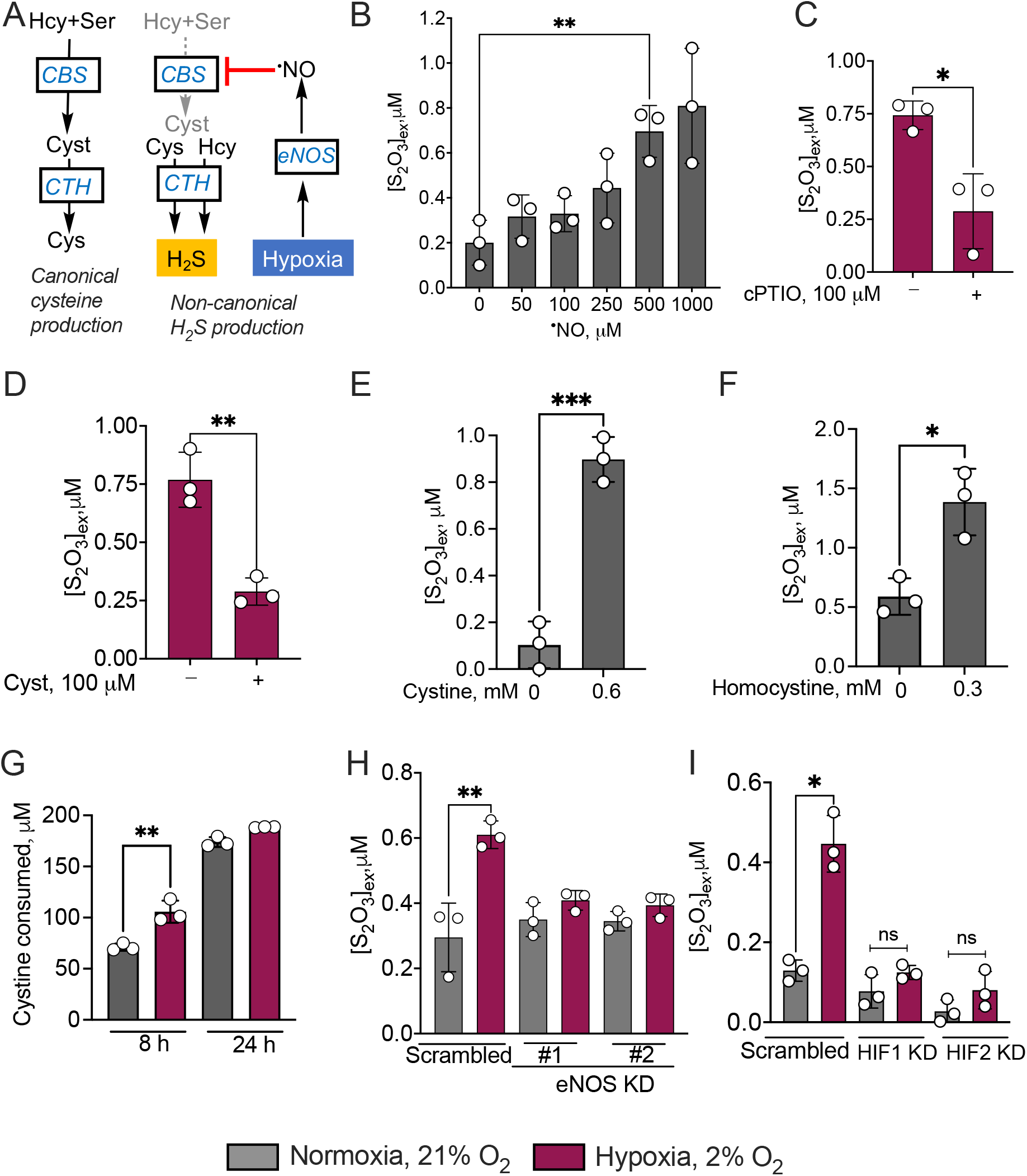
Intersection of •NO and HIF signaling in hypoxic regulation of H_2_S homeostasis. **(A)** Scheme showing •NO-induced switching at CTH from cysteine- (*left*) to H_2_S-generation (*right*). **(B)** Concentration-dependent increase in extracellular thiosulfate [S_2_O_3_^2-^]_ex_ accumulation by DETA NONOate in EA.hy926 cells under normoxia (4 h, n=3, **p=0.005). **(C, D)** Hypoxic thiosulfate accumulation after 24 h is decreased by the •NO scavenger cPTIO (C) (n=3, *p=0.014) and by cystathionine (D, n=3, **p=0.003). **(E,F)** Cystine and homocystine increase [S_2_O_3_^2-^]_ex_ at 24 h in EA.hy926 cells (n=3 ***p=0.0006 and *p=0.013). (**G**) EA.hy926 scrambled controls show faster extracellular cystine consumption at 8 h (n=3 **p=0.006) **(H,I)** Knockdown of eNOS (H, using shRNA sequences 1 and 2, n=3, **p=0.008) or HIF1 or HIF2 (I, n=3, *p=0.002), in EA.hy926 cells decreased hypoxic [S_2_O_3_^2-^]_ex_ accumulation at 24 h. Error bars represent ± S.D.

### SQOR is needed for hypoxic endothelial cell proliferation

SQOR but not ETHE1 deficiency (Figure S4A and B) attenuated Ea.hy936 cell proliferation, which profoundly exacerbated under hypoxia (Figure 3A,B). SQOR KD in a second endothelial cell line HMEC-1, also lowered cell proliferation (Figure S4C and S5A). The vasculogenic potential for forming capillary like structures as monitored by the tube formation assay, revealed that SQOR KD were defective under both normoxic and hypoxic conditions (Figure 3C,D and Figure S5B,C). SQOR KD in EA.hy926 and HMEC-1 cells diminished colony formation (Figure S5D-G). VEGF-induced endothelial cell angiogenic sprouting assessed with a microphysiologic platform^36^, revealed significantly lower tip cell formation of SQOR KD cells under normoxia (Figure 3E,F). SQOR deficiency also decreased cell proliferation as measured by EdU labelling in normoxia (Figure 3E,G). These data suggest that SQOR deficiency compromises VEGF-induced angiogenic sprouting in endothelial cells.

**Figure 3.**
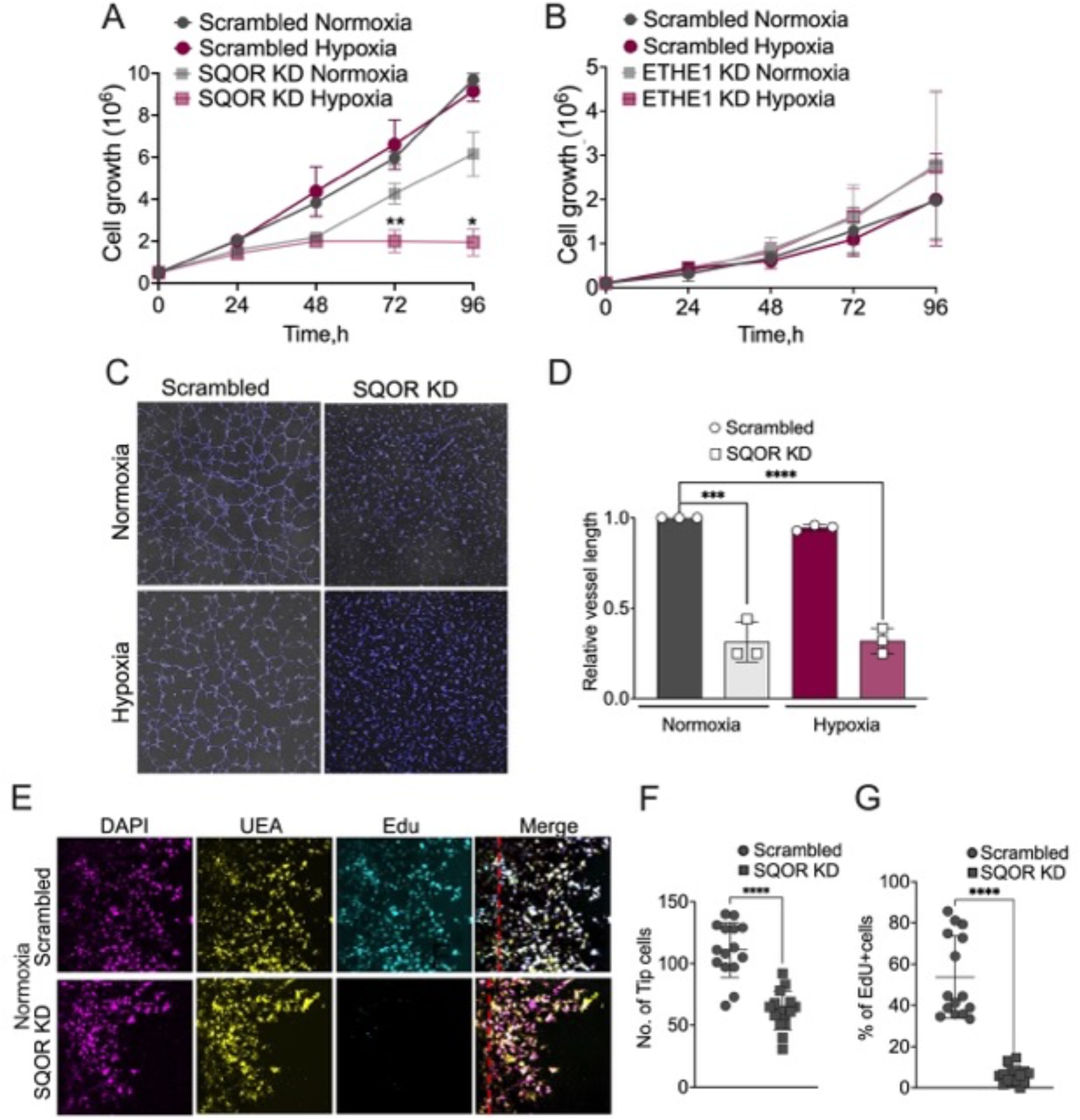
SQOR knockdown decreases endothelial cell proliferation. **(A, B)** SQOR KD decreases proliferation of EA.hy926 cells, which is more pronounced under hypoxic conditions (*p=0.0470,**p=0.0056) (A), while ETHE1 KD has no effect (B). (**C, D**) SQOR KD leads to defective tube formation (C) as evidenced by decreased vessel length (D). n=3, ***p=0.0004, ****p<0.0001. (**E,F,G**) Representative images for VEGF (10 ng/ml) induced sprouting of Ea.hy926 cells (E), quantitation of the number of tip cells (F,****p<0.0001) and Edu+ proliferating cells (G, p<0.0001). The dashed red line in E indicates the edge of the parent vessel. n=15 (F and G) and error bars are ± S.D.

### Reductive shift in the mitochondrial NADH pool limits SQOR-deficient endothelial cell proliferation

SQOR deficiency is predicted to impair H_2_S oxidation, which was confirmed by the reduced rate of H_2_S clearance and thiosulfate accumulation by SQOR KD versus scrambled controls (Figure 4A,B). The expected 2:1 stoichiometry between H_2_S consumed and thiosulfate produced was seen in control but not SQOR KD cells, where thiosulfate levels were comparable to the background levels. Due to its volatility, a sizeable abiotic loss of H_2_S was seen even in the absence of cells.

**Figure 4.**
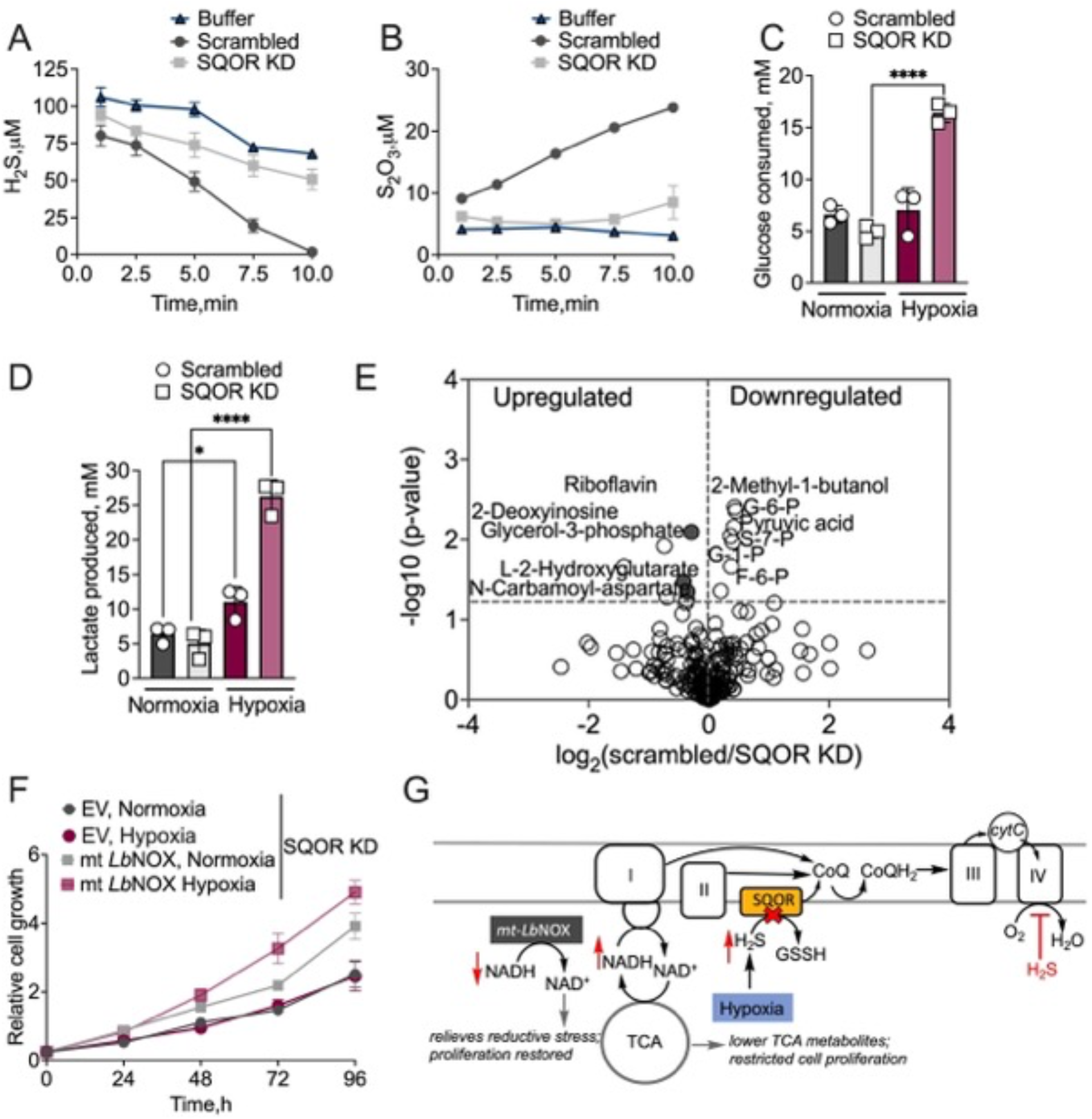
Reductive shift in the mitochondrial NADH pool limits endothelial cell proliferation. **(A,B)** Kinetics of H_2_S consumption (A) and thiosulfate production (B) by 10% (w/v) EA.hy926 cell suspensions (n=3) revealed a 2:1 H_2_S:S_2_O_3_^2-^ ratio for scrambled but not SQOR KD cells. (**C, D**) Comparison of glucose consumption (n=3, ****p<0.0001) (C), and lactate production (n=3 *p=0.0345, ****p<0.0001) (D) by Ea.hy926 grown for 48 h in 2 versus 21% O_2_. (**E**) Metabolomics data reveal increased levels of NADH-sensitive metabolites under normoxia in SQOR KD cells. (**F**) Expression of mt-*Lb*NOX alleviates the growth restriction of SQOR KD EA.hy926 cells under normoxic and hypoxic conditions. Error bars are ± S.D. (**G**) Model explaining how SQOR KD exacerbates reductive stress due to increased hypoxic H_2_S synthesis, which can be alleviated by mt*Lb*NOX.

SQOR KD cells exhibited increased glucose consumption under hypoxia, which was paralleled by extracellular lactate accumulation (Figure 4C,D). Earlier studies reported K_ATP_ channel activation as a mechanism of H_2_S regulation of endothelial cell proliferation^8^. However, neither the K_ATP_ channel activator SG209 nor the channel blocker, glybenclamide, affected proliferation of scrambled or SQOR KD EA.hy926 or HMEC-1 cells (Figure S5H,I). Metabolomics analysis of normoxically grown SQOR KD versus scrambled EA.hy926 cells revealed accumulation of glycerol-3-phosphate, L-2-hydroxyglutarate and N-carbamoyl asparate (Figure 4E, Table S2). These changes suggested the hypothesis that SQOR deficiency leads to a reductive shift in the mitochondrial pyridine nucleotide pool as previously seen in H_2_S treated cells^23^. To test this hypothesis, mito-*Lb*NOX, a water-forming NADH oxidase targeted to the mitochondrion^37^, was expressed in the background of SQOR knockdown in Ea.hy926 cells (Figure S5J). Mito-*Lb*NOX promoted growth of SQOR KD cells under both hypoxia and normoxia (Figure 4F). These data support the model that hypoxic activation of H_2_S synthesis requires SQOR activity to avert complex IV inhibition and ETC backup, and to activate glycolysis for ATP generation while the TCA cycle is used for macromolecular precursor synthesis (Figure 4G).

### Tumor growth and angiogenesis require SQOR

We tested the physiological relevance of SQOR deficiency on angiogenesis in a tumor xenograft model in which syngeneic YUMM5.2 mouse melanoma cells were transplanted subcutaneously in 8-10 week old control (*Sqrdl* ^fl/fl^) or SQOR KO (*WB^cre^ Sqrdl^fl/fl^*) mice (Figure 5A). Whole body Cre recombinase was induced by daily intraperitoneal tamoxifen injections for 5 days and validated in colon and liver by Western blot analysis as well as by the concentration of urinary thiosulfate, a marker of H_2_S oxidation, 10 days post tamoxifen injection (Figure 5B,C). Tumors implanted in SQOR KO mice showed significant reduction in mass compared to littermate wildtype mice (Figure 5D,E) and a marked decrease in tumor Ki67 staining (Figure 5F,G). Immunohistochemical analysis of CD31, an endothelial marker, revealed significantly decreased vessel number (Figure 5H,I), consistent with the *in vitro* data (Figure 3D,F,G). In contrast, histological examination of lung sections revealed no difference in CD31 staining (Figure S6), indicating that blood vessel maintenance was not impacted following induction of SQOR deficiency at least over the time frame of the experiment. These results highlight the importance of SQOR in neovascularization.

**Figure 5.**
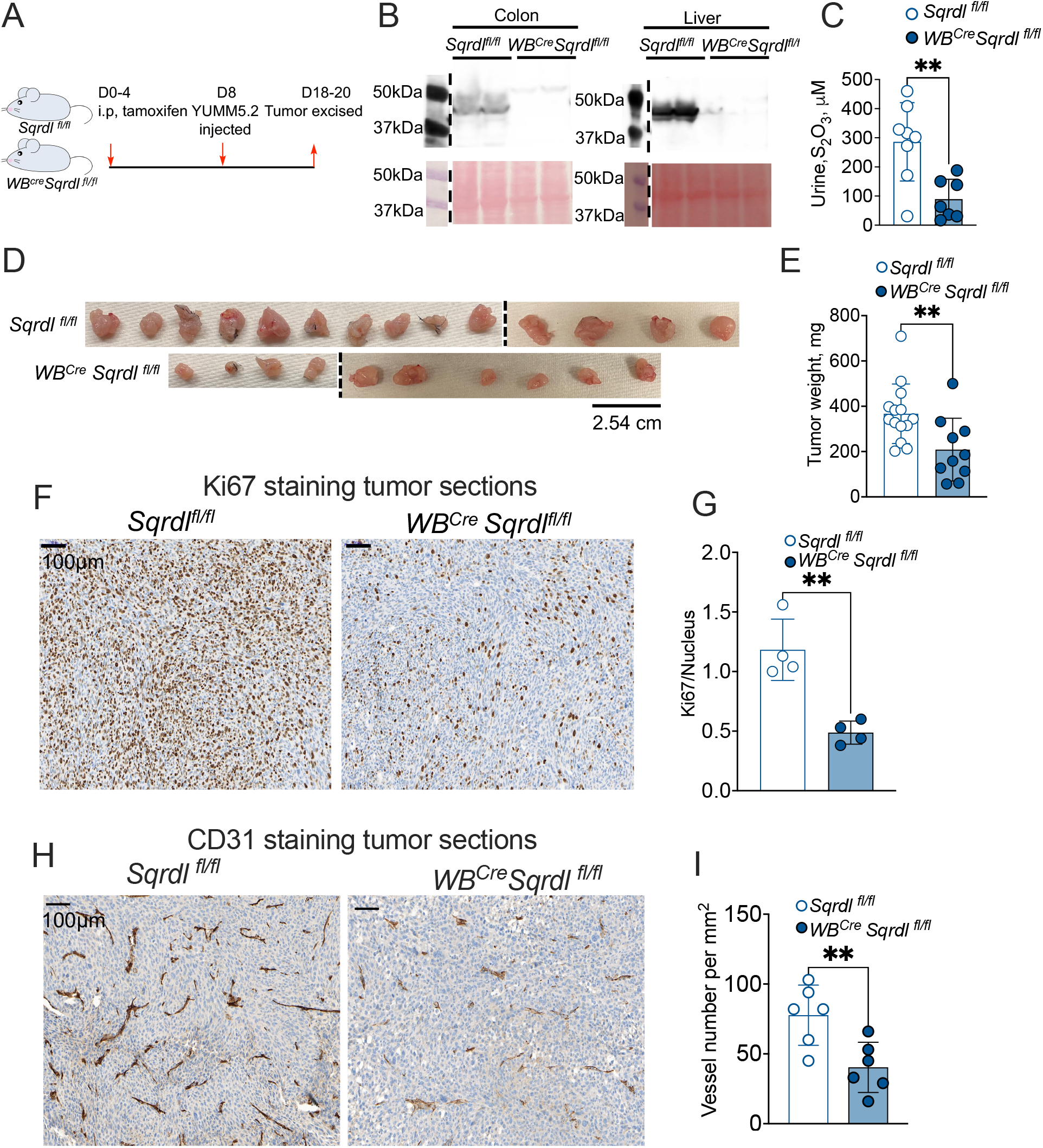
SQOR activity supports tumor growth and angiogenesis. **(A)** Scheme showing induction of SQOR KO and syngeneic tumor generation. **(B)** Western blot analysis validating SQOR KO in colon and liver of WB^Cre^ Sqrd^lfl/fl^ mice. **(C)** Urinary thiosulfate is lower in WB^Cre^ Sqrd^lfl/fl^ versus controls. **(D)** Tumors harvested from WB^Cre^ Sqrd^fl/fl^ (n=10) versus Sqrd^lfl/fl^ (n=14) mice. **(E)** Comparison of tumor mass in the two groups (**p=0.0093). **(F,G)** Representative Ki67 staining of tumor sections (F) and quantitation (G) (n=4 mice from each group, **p=0.0023). **(H,I)** Representative CD31 staining of tumor sections (H) and quantitation (I) (n= 3 mice from each group, two different areas, **p=0.0086). Error bars are ± S.D.

### SQOR deficiency lowers muscle angiogenesis in hind limb ischemia mice model

Femoral artery ligation induces ischemia and promotes angiogenesis in the distal part of the limb and especially in the gastrocnemius muscles. Surgery was performed 2 weeks after tamoxifen injection (Figure 6A). Blood flow was similarly impaired in both control and KO mice after day 1 post-surgery establishing the validity of the model. Neovascularization, which is indicated by the return of the blood flow determined by Doppler flow imaging, was lower in SQOR KO mice compared to controls on day 3 (Figure 6 B,C,D). By day 8, significant differences in the blood flow were not observed, presumably due to artegiogeneis (Figure S7A,B). However, CD31 staining of gastrocnemius muscle sections on day 9 revealed lower capillary to muscle fiber ratio in WB^Cre^*Sqrdl*^fl/fl^ mice versus the *Sqrdl*^fl/fl^ controls (Figure 6E,F). These data are consistent with a role for SQOR in ischemia induced neovascularization.

**Figure 6.**
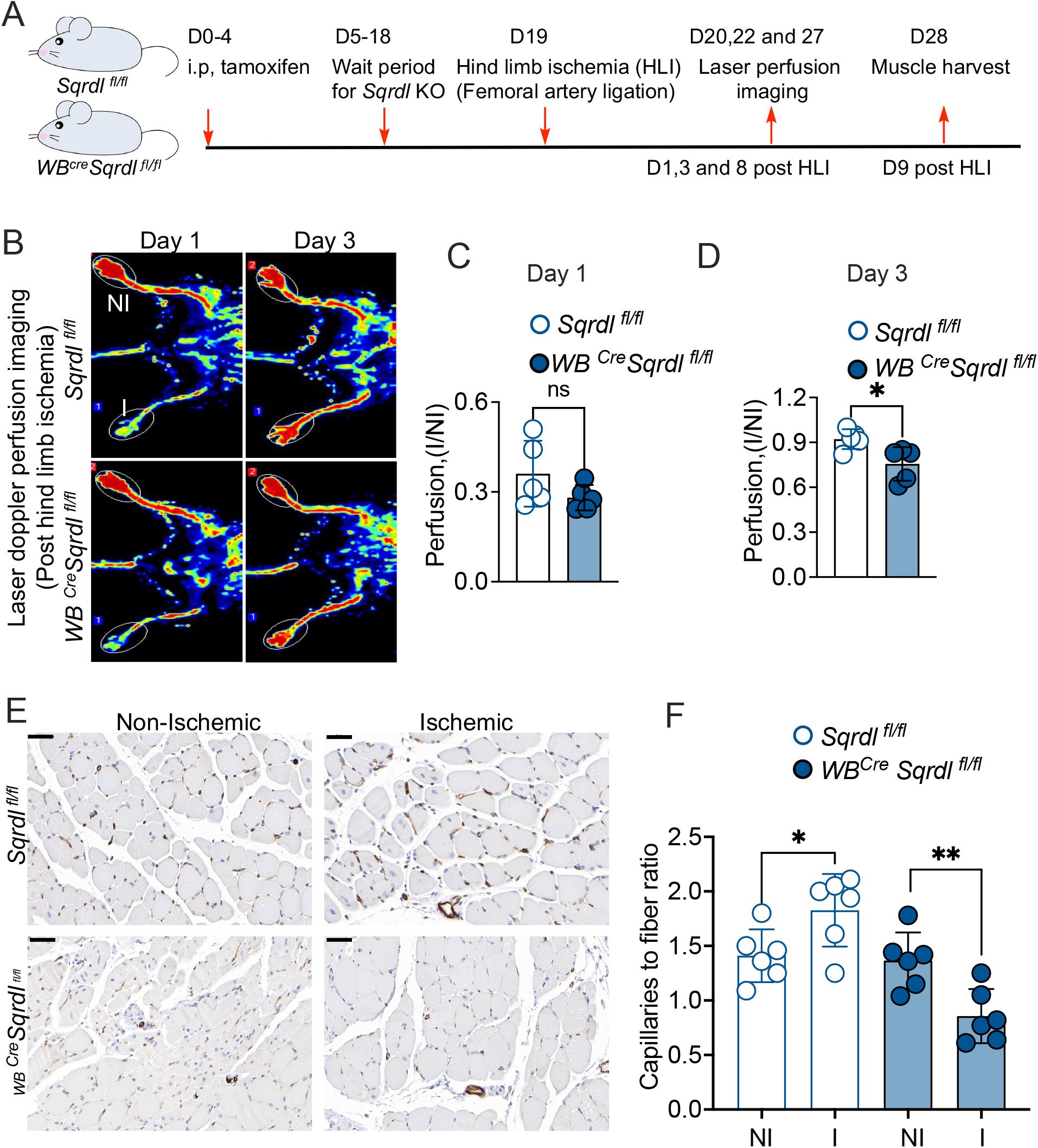
Loss of SQOR decreases angiogenesis in a hind limb ischemia model. (**A**) Experimental design for induction of hind limb ischemia by femoral artery ligation in SQOR KD versus control mice. (**B**) Laser doppler perfusion imaging at days 1 and 3 post hind limb ischemia. The top non-ischemic (NI) and bottom ischemic (I) limbs are labeled. The white ovals outline the distal regions where angiogenesis is stimulated following ischemia. (**C** and **D**) Blood perfusion is plotted as the ratio of ischemic:non-ischemic limb. (**E**) CD31 staining of gastrocnemius muscle harvested on day 9. Scale bar represents 50 μm. (**F**) Quantitation of CD31 staining as the ratio of capillaries:fiber from 6 different areas in 4 mice. Error bars are ± S.D.

## Discussion

While the interdependence of •NO and H_2_S on vascular function is known, the molecular basis by which eNOS deficiency inhibits H_2_S-mediated neovascularization or how CTH loss lowers sprout formation induced by the •NO donor, 2-(N,N-diethylamino)-diazenolate-2-oxide, has remained elusive^7^. In this study, we demonstrate that the hypoxic induction of eNOS disconnects metabolic flux between CBS and CTH in the canonical transsulfuration pathway (Figure 2A), decreases GSH synthesis, and promotes H_2_S metabolism. Hypoxic-enhancement of extracellular cystine consumption presumably provisions CTH with the substrate for H_2_S production. In this model, •NO acts as an upstream regulator of H_2_S biogenesis by promoting hypoxic H_2_S synthesis instead of the canonical cysteine synthesis.

Previous studies have linked the hypoxic lowering of GSH to its utilization in quenching reactive oxygen species (ROS)^38,39^. We demonstrate that endothelial cells prioritize cysteine utilization for H_2_S generation over GSH production, leading to a decreased GSH pool size. The oxidative pentose phosphate pathway is a major source of NADPH and increased turnover of oxidized GSH by glutathione reductase might be involved in protection against ROS under hypoxia. In this context, inhibition of the oxidative pentose phosphate pathway has been shown to decrease HUVEC viability and impair blood vessel maturation^40,41^.

The hypoxia-HIF axis regulates both tip cell migration and stalk cell proliferation through transcription of pro-angiogenic factors including VEGF and NOS^42^. Our results indicate that upregulation of H_2_S metabolism lies downstream of HIF stabilization since HIF1 or HIF2 KD cells under hypoxia decreased, while FG4592 treated cells under normoxia increased, H_2_S oxidation as monitored by the proxy marker thiosulfate (Figures 2I and S3D). Even highly glycolytic cells need a functional TCA cycle to generate macromolecular precursors as evidenced by inhibition of endothelial cell proliferation by ablation of complex III^26,43^. A balance between endothelial cell migration and proliferation is key to angiogenesis^36^. While the ETC inhibitors, cyanide, phenformin and oligomycin stimulate migration^9^, H_2_S oxidation also regulates proliferation (Figure 3A). Furthermore, a role for H_2_S *per se* rather than a downstream reactive sulfur species in regulating proliferation is indicated by the absence of an observable effect in ETHE1 KD cells (Figure 3B). Since ETHE1 oxidizes the SQOR product, glutathione persulfide, to sulfite^44^ (Figure 1A), this reactive low molecular weight persulfide accumulates in ETHE1 KD cells, triggering formation of additional poly- and persulfide derivatives^45^. We posit that H_2_S exerts its pro-angiogenic effect by stimulating SQOR activity and inhibiting complex IV, which shifts energy metabolism from oxidative phosphorylation toward aerobic glycolysis^9,23,25^. This shift would support rapid ATP production to meet the energy demands of motile tip cells.

The efficiency of the sulfide oxidation machinery is important for maintaining very low steadystate levels of sulfide^21,46^. A spike in endogenous H_2_S levels^29,47^ could synergize with hypoxia to dampen ETC flux by increasing the *K*_M_ for O_2_ of cytochrome C oxidase, which is very low (<0.2 to 1 μM, corresponding to <0.02–0.1% O_2_ tension)^22,48^. Thus, even in the colon where the estimated O_2_ tension is as low as 0.4%, the O_2_ concentration is 4 to 20-fold higher than the *K*_M_ for cytochrome C oxidase, and should be insensitive to hypoxic regulation^49^. Our study helps explain this apparent paradox, by demonstrating that while endothelial cells proliferate equally well under normoxic versus hypoxic conditions, SQOR deficiency drastically reduces proliferation particularly under hypoxia (Figure 3A). We interpret this data as evidence for a steady-state increase in H_2_S levels due to SQOR deficiency with a concomitant and sustained inhibition of the ETC (*K*_i_(H_2_S) ~200 nM^50^), which in turn, induces reductive stress and limits TCA cycle activity, inhibiting proliferation (Figure 4H). Consistent with this model, hypoxically grown SQOR-deficient endothelial cells accumulate 2-hydroxyglutarate, a product of a-ketoglutarate reduction and a marker of ETC dysfunction^51,52^. Additionally, glycerol-3 phosphate (G3P) and N-carbamoyl aspartate also increase under these conditions. G3P shuttles reducing equivalents generated by cytosolic G3P dehydrogenase during the NADH-dependent oxidation of dihyroxyacetone phosphate. The mitochondrial isoform of the same enzyme then transfers electrons transiently to FAD as it catalyzes the oxidation of G3P to dihydroacetone phosphate and uses CoQ as the final electron acceptor. N-carbamoyl asparate is a precursor of dihydroorotate in the pyrimidine biosynthesis pathway. Accumulation of G3P and carbamoyl phosphate is consistent with a reductive shift in the ETC. Thus, while H_2_S modulates the cellular response to hypoxia in endothelial cells, its sustained accumulation in SQOR deficient cells is detrimental for proliferation.

The physiological relevance of SQOR in angiogenesis is exemplified by decreased tumor angiogenesis in SQOR KO mice, suggesting that H_2_S build up due to induction of intratumoral H_2_S synthesis in the hypoxic tumor microenvironment, is sufficient to restrict tumor growth. Similarly, in the femoral ligation model for hind limb ischemia, SQOR KO mice demonstrated reduced blood flow as well as decreased muscle angiogenesis compared to controls. A previous *in vitro* study on SQOR knockdown in Hepa 1-6 cells identified a protective role for H_2_S oxidation in ischemia/reperfusion injury ^53^. Other studies using murine SQOR deficiency models induced either by global SQOR KO or mislocalization via deletion of the mitochondrial leader sequence, reported that the homozygous SQOR^-/-^ pups failed to thrive and did not survive past 8 or 10 weeks of birth^29,54^. The effect of SQOR deficiency on the development of blood vessels was not evaluated in these studies. In our study, whole-body SQOR KO was induced ~2 months after birth and the absence of an effect in lung CD31 expression indicates that SQOR is not required for blood vessel maintenance (Figure S6). Our model thus allowed evaluation of the effects of SQOR deficiency on neovascularization in pathological contexts in adult mice.

The interdependence of O_2_ and H_2_S metabolism in ETC flux has significant physiological relevance as evidenced by the sensitivity of the brain to ischemia due to low levels of SQOR^29^, further demonstrating that low H_2_S clearance capacity represents a metabolic vulnerability. We have previously demonstrated that sustained inhibition of the ETC by H_2_S leads to recruitment of a redox cycle between SQOR and complex II, which prioritizes H_2_S removal and CoQ regeneration, using fumarate as a terminal electron acceptor^27^. While hypoxia also leads to utilization of fumarate as a terminal electron acceptor in multiple tissues^28^, the contribution of this redox cycle in endothelial cells and its modulation by H_2_S remain to be assessed.

In summary, we demonstrate that hypoxic reprogramming of the transsulfuration pathway by •NO induces H_2_S in endothelial cells and is crtical for angiogenesis. The study revealed the underlying mechanism by which •NO, H_2_S and O_2_ metabolism intersect to regulate ETC redox state and angiogenesis. Finally, our study revealed a key role for sulfide oxidation rather than reactive persulfide species in angiogenesis and nominated SQOR as a rational drug target for anti-angiogenesis therapy in different disease settings.

## Acknowledgements

This work was supported in part by the grants from the National Institutes of Health (GM130183 to RB, R01CA248160 to CAL and R01CA148828 and R01CA245546 to YS), the American Heart Association (826245 to RK), the American Physiology Society and Crohn’s and Colitis Foundation (019463 and 1003279 to RS) and the National Institute of Dental and Cranofacial Research (T32DE00705745 to HLH). We acknowledge Aaron Landry and Wesley Huang (University of Michigan) for their technical help with generating the SQOR KD in Ea.hy926 cells and with harvesting murine samples, respectively. We acknowledge physiology phenotyping core at University of Michigan manager Steven Whitesall for hind limb ischemia surgery and laser doppler perfusion imaging. We thank Michael Mattea from University of Michigan centre for gastrointestinal research for histology studies.

## Author contributions

RK, YS and RB conceptualized the study and RK performed and analyzed the majority of the experiments and was assisted by: VV-[^35^S]-Met flux and glucose consumption assays, PS-proliferation assays, HB-tumor xenograft experiment. HH and BB-performed the tip sprouting assay, RS-HIF1/2 Western blots and YUMM 5.2 culture, AA and CAL-metabolomics data generation and analysis. RK and RB drafted the manuscript and all authors edited and approved the final version.

## Materials

Sodium sulfide nonahydrate (431648), sodium sulfite (S0505), sodium thiosulfate (217263), L-glutathione reduced (G4251), L-glutathione oxidized (G4501), L-cystine dihydrochloride (C6727), L-cysteine hydrochloride (C6852), L-arginine monohydrochloride (A5131), doxycycline (D3447), puromycin (P8833), protease inhibitor cocktail for mammalian tissue extract (P8340), RIPA lysis buffer (R0278), cystathionine (C3633), D-glucose (G7021), dimethyl sulfoxide (D2650), uridine (U3003), hydrocortisone (H0888), and glybenclamide (G0639) were from Sigma. RPMI 1640 (11875-093), DMEM (11995-065), fetal bovine serum (10437-028), trypsin-EDTA (25300-054), penicillin-streptomycin (15140-122), geneticin (10131-035),epidermal growth factor (PHG0311), PBS (10010-023), DPBS (14040-133), MCDB 131 (10372-019), sodium pyruvate (11360-070), and trypan blue (15250-061) were from Gibco. Vascular cell basal medium (PCS 100-030), Dulbecco phosphate buffered saline (30-2200), endothelial cell growth kit (PCS 100-041), trypsin EDTA (PCS 999-03), and trypsin neutralizing solution (PCS 999-04) were from ATCC. Carboxy PTIO (81540), DETA NONOate (82120), and FG4592 were from Cayman chemical company. L- [^35^S]-Methionine (NEG009A500UC) was from Pekin Elmer. eNOS (9572S, Cell Signaling Technology), Ki67 (12202T, Cell Signaling Technology), HIF-1α (ab179483, Abcam), HIF-2α (A700-003, Bethyl Lab), CD31 (28083-1AP, Proteintech) SQOR (17256-1AP, Proteintech), and Flag (F1804, Sigma) antibodies and the secondary anti-rabbit horseradish peroxidase-linked IgG antibody (NA944V, GE Healthcare for western blotting and 7074S Cell signaling Technology for IHC) were purchased from the indicated vendors. UEA DyLight 649 labeled Ulex europaues Agglutinin-1 (UEA, 1:200, Vector Labs), EdU (ClickIT EdU, Life Technologies), tamoxifen (10540-29-1, MCE), formalin (Fisher Scientific SF98-4), SG-209 (T24786, TargetMol), EZ prep solution (950-102), DISCOVERY Cell conditioning 1 buffer- 950-500, Reaction buffer concentrate-950300, omniMap anti-Rb-HRP- 760-4311, ChromoMap DAB- 760-159 from Ventana Medical Systems (Roche diagnostics) and hematoxylin stain, Gill II (newcomersupply) were from the indicated vendors.

## Online Methods

### Cell culture

EA.hy926 and HEK293 were cultured in Dulbecco’s modified eagle medium (DMEM). HMEC-1 (human dermal microvascular endothelial cells) were cultured in MCDB131 with 10 ng/mL epidermal growth factor, 1 μg/mL hydrocortisone and 10 mM glutamine. HUVEC (human umbilical vein endothelial cells) were cultured in vascular cell basal medium to which the endothelial cell growth kit was added to obtain final concentration of the following growth factors: VEGF (5 ng/mL), rhEGF (5 ng/mL), rhFGF basic (5 ng/mL), rhIGF-1 (5 ng/mL), L-glutamine (10 mM), heparin sulfate (0.75 units/mL), hydrocortisone hemisuccinate (1 μg/mL), and ascorbic acid (50 μg/mL). Cell lines were cultured in the specified medium: HepG2 (Eagle minimum essential medium containing 2 mM glutamine), 143B (DMEM supplemented with100 μg/mL uridine) and SH-SY5Y cells (DMEM/F12). All culture media were supplemented with 1% penicillin-streptomycin and 10% FBS except the medium for HUVECs, which contained 2% FBS. Cells were cutured in 5% CO_2_ incubators with humidified ambient air containing 21% O_2_ (normoxia) or with 93% N_2_, 5% CO_2_ and 2% O_2_ (hypoxia chamber).

### Mice

All experiments were performed with the approval from Institutional Animal Care and Use Committee (IACUC) at the University of Michigan. Animals were maintained under standard housing conditions with ad libitum access to food and water and 12h light-day cycle.

### SQOR^fl/fl^ knockout mice generation

Sqrdl^loxP/+^ mice were generated by Cyagen Biosciences (Santa Clara, CA). Briefly, the targeting vector was constructed by inserting one SDA (self-deletion anchor)-flanked neomycin cassette and two loxP sites flanking exon 7 of *sqrdl* and then electroporated into embryonic stem (ES) cells from C57BL/6N mice. The transfected ES cells were subject to G418 selection (200 μg/mL), 24 h post electroporation. G418 resistant clones were picked and amplified in 96-well plates. The following primers were used for screening targeted clones: F:5’-AGTTTCAAGGCTCTAAACATCTCCT-3’ R:5’-AATACCTTCAATAGGAGAGATGGGG-3’. The positive clones were expanded and further characterized by Southern blot analysis, injected into C57BL/6 embryos and reimplanted in CD1 pseudo-pregnant females. Following one subsequent cross with C57BL/6 animals, the Neo transgene was removed, and the *sqrdl*^loxP/+^ mice were generated. The following primers used to identify neo deletion (F1:5’-TTTTCTTCTGCCTAAAACCCTGC-3’ R1:5’-AATCTAAAAGGCAATTCTCCCCATC-3’’). An inducible whole body disruption of *sqrdl* was generated by crossing Sqrdl^loxP/loxP^ mice with CAGG-CreER mice (Jax # 004682).

### [^35^S]-Methionine flux into GSH

Cells were seeded in 6 cm plate at a density of 2.4 million cells except HUVEC (1.2 million/plate plate) and allowed to settle overnight (15-18h), washed twice with 2 mLof 1X PBS pH 7.4 before 4 mL of fresh medium was added except for HUVEC cultures, in which (2 mL medium/6 cm plate) was added. Each culture medium also contained either 5 μCi/mL [^35^S]-methionine (EA.hy926, HMEC-1 and HUVEC) or 2.5 μCi/mL [^35^S]-methionine and cells were incubated for 24 h under normoxic or hypoxic conditions except for HUVECs, which were incubated for 16 h. Then, the medium was aspirated, the cells were washed twice with 1X PBS and scraped in 150 μL 1X PBS to obtain cell suspensions. For the analysis of radiolabel incorporation in GSH, 100 μL of each cell suspension was mixed with an equal volume of metaphosphoric acid solution (135 mM metaphosphoric acid, 5 mM EDTA, and 150 mM NaCl), vortexed and stored at −20 °C. For protein quantitation, 30 μL of cell suspension was mixed with an equal volume of RIPA buffer with protease inhibitor and stored at −20 °C. The protein concentration in thecell suspension was measured using the Bradford reagent.

To measure radiolabel incorporation into GSH, cell samples were treated and analyzed by HPLC as described previously^55,56^. Briefly, samples were thawed, vortexed and centrifuged at 12,000 x g for 5 min at 4° C. The supernatant (150 μL) was mixed with 15 μL of iodoacetic acid (14 mg/mL) to alkylate the thiols, the pH was adjusted (using a pH paper) to 7–8 with saturated potassium carbonate using a pH paper, and the mixture was incubated for 1 h at room temperature in the dark. Then, an equal volume of 2,3-dinitrofluorobenzene (1.5 % v/v in absolute ethanol) was added to derivatize amino groups and incubated at room temperature for 4 h in the dark. Cystine, cysteine, and GSH were separated by HPLC using a Bondapak NH_2_ column (Waters) column (300 mm x 3.9 mm, 10 μm,Waters) at a flow rate of 1 mL/min. The mobile phase consisted of solution A (80% methanol in water) and solution B which was prepared by mixing 154.9 g of ammonium acetate in 100 ml of water and 400 ml of glacial acetic acid and adding 150 ml of resulting solution to 300 ml of solvent A. The mobile phase comprised: 0 to 10 min, isocratic 25% solution B; 10 to 30 min, linear gradient from 25 to 100% solvent B. Elution of metabolites was monitored by absorbance at 355 nm. The column was equilibrated with 30% solvent B before samples (50-100 μl) were injected. Peak elution was monitored by absorbance at 355 nm. The concentration of metabolites (cystine, cysteine and GSH) was determined by comparing the integrated peak area with a calibration curve generated for each compound. Incorporation of [^35^S]-methionine into GSH was determined by measuring the radioactivity in the corresponding chromatographic fraction and the data were normalized to protein concentration.

### Extracellular thiosulfate quantitation

Cells were seeded similarly as described above and the following day, cells were washed twice with 2 mL 1X PBS and switched to fresh medium (3 mL) and cultured at 2 or 20% O_2_. Extracellular thiosulfate was determined in the conditioned medium at 24 h (and in some cases 4 h). The medium from HUVECs cultured in DMEM was harvested at 16 h. Thiosulfate levels were quantified in each culture medium to obtain blank values. For thiosulfate derivatization, 45 μL of culture medium was mixed with 2.5 μL of Tris base (1M) and 2.5 μL monobromobimane (60 mM). The mixture was vortexed and incubated at the room temperature in the dark for 10 min. Next, proteins in the derivatized mixture were precipitated using metaphosphoric acid (16.8 mg/mL). Samples were centrifuged at 12,000 x g for 5 min at 4° C and the supernatant was collected in the dark and stored at −20 °C till further use.

Zorbax Eclipse XDB-C18 column (5 μm, 4.6 × 150 mm, Agilent) was used to separate thiosulfate using ammonium acetate/methanol buffer system with the gradient as follows-Solution A contained 100 mm ammonium acetate, pH 4.75, and 10% methanol; Solution B contained 100 mm ammonium acetate, pH 4.75, and 90% methanol. The percentage of B was increased in the gradient as follows: 0–10 min, linear 0 to 20%; 10–15 min, linear 20 to 50%; 15–j20 min, isocratic 50%; 20–22 min, 50 to 100%; 22–27 min, isocratic 100%; 27–29 min, linear 100 to 0%; 29–35 min, isocratic 0%. Thiosulfate was detected using excitation and emission wavelengths set at 390 nm and 490 nm, respectively. The column was calibrated with known concentrations of derivatized sodium sulfide, sodium sulfite, and sodium thiosulfate.

### *shRNA knockdowns and Lb*NOX *expression*

The pLKO.1 vector containing shRNAs were purchased from Sigma. The following clone IDs were used for SQOR (TRCN0000039004, TRCN0000039006), ETHE1 (TRCN0000083454, TRCN0000083455), eNOS (TRCN0000045476, TRCN0000045477, TRCN0000414816). pZIP-HIF1 and HIF2 was from the Shah lab and the pINDUCER empty vector or containing mitochondrial *Lb*NOX was from the Lyssiotis (University of Michigan). shRNA (2.5 μg) containing plasmids were submitted to the University of Michigan Vector Core for lentiviral packaging. To generate shRNA induced knockdowns, cells were seeded at a density of 7.5 X 10^4^ in a 6-well plate containing 2 ml culture medium per well. The next day, the medium was changed and cells were transduced with the optimized viral titer for 48 h in the presence of polybrene (8μg/mL). The medium was then replaced with virus free medium and growth was continued for a further 24 h. Puromycin (1 μg/mL) was used to select knockdown cells while geneticin (500 μg/mL) was used for the selection of mito-*Lb*NOX-expressing cells. A combination of puromycin (1 μg/mL) and geneticin (500 μg/mL) was used for *Lb*NOX expression in the SQOR KD background. Once the wells were confluent, they were transferred to 10 cm plates. Selection was continued for ~2 weeks and the knockdown efficiency and *Lb*NOX expression were estimated by Western blotting.

### Western blotting

Three 10 cm plates with confluent EA.hy926 cells were washed twice with 1X PBS, scraped in 750 μL 1X PBS and pooled. After centrifugation at 1700 x g, 5 min at 4 °C, the supernatant was discarded, and the pellet was dissolved in 100 μL of RIPA buffer containing protease inhibitor and vortexed. Three freeze-thaw cycles and vortexing resulted in efficient cell lysis and the supernatant was collected after centrifugation at 12,000 x *g*, 5 min at 4° C. Then, the lysate was mixed with 4X sample buffer to obtain final concentration of 2 μg/μl, heated at 95° C for 5 min and used immediately for Western blot analysis or stored at −80 °C for up to 2 weeks. Protein was separated on a 10-12% SDS polyacrylamide gel and transferred to a PVDF membrane. Membranes were incubated overnight at 4 °C with primary antibodies with the following dilution SQOR (1:2000), ETHE1 (1:1000), eNOS (1:2000), HIF1 (1:1000), HIF2 (1:1000) anti-Flag (1:2000). The secondary antibody, horseradish peroxidase–linked anti-rabbit IgG was used at a 1:10,000 dilution. Blots were developed with KwikQuant Digital-ECL substrate (KwikQuant), and images were collected with a KwikQuant Imager. Equal loading was verified by Ponceau S staining of membranes or with actin for HIF1 and HIF2.

### Proliferation assay

Cells were seeded at a density of 10,000-50,000 in 2 mL of media and grown overnight in 6 well plates under standard cell culture conditions (37 °C and 5% CO_2_ and ambient air). The medium was changed (2 mL) the next day, and cells were placed in 21% or 2% O_2_ incubators. Cells were counted starting 24 h later and every 24 h for continued for 4 days. For this, cells were washed twice with 2 mL 1X PBS, and 500 μL 0.05% trypsin (0.25% for HMEC-1) was added and placed at 37 °C for 5-7 min. Then, the cells were resuspended with 500 μL of the medium and centrifuged at 4°C for 5 min. The supernatant was discarded and the cell pellet was resuspended in 50 μL of the medium (as specified for each cell line above) and diluted 1:1 (v/v) with trypan blue and 20 μL of the resulting sample was counted using a Cellometer (Nexelcom).

### Clonogenic assay

Cells were seeded ina 6-well plate at a density of 500 cells per well with each containing 2 ml of culture medium, and grown overnight under standard cell culture conditions (37 °C and 5% CO_2_ and 21% O_2_). The next day, cells were placed in 2 or 21% O_2_ incubators and grown for 10-12 days with fresh medium change every 3-4 day. The fresh medium for cells growing in 2% O_2_ was placed in the hypoxic chamber overnight prior to use. Once colonies were visible, the medium was aspirated and washed twice with 1X PBS, fixed with 10% buffered formalin for 20 min and then incubated in crystal violet solution (0.5% crystal violet in 20% methanol) for 35 min. Then, the wells were washed 6 times with 4 ml of distilled water to remove residual stain, dried in an inverted position overnight and imaged. Colonies were counted for quantitation and any colony containing less than 5 pixels was not included in the analysis.

### Tube formation assay

Corning matrigel was thawed on ice and 200-250 μl was added per well in a 24-well plate using chilled tips and placed in a 37 °C incubator for 30 min to coat the wells. 1.2X10^5^ EA.hy926 and HMEC-1 (scrambled and SQOR knockdown cells) were seeded in DMEM (160 μl) with 10% FBS and 1% penicillin/streptomycin and diluted 1:1 with incomplete DMEM (without FBS, 160 μl) to obtain a final FBS concentration of 5% and placed in a 20% O_2_ incubator for 1 h before being switched to a 2% incubator. Cells were imaged with BioTek, BioSpa Live Cell Analysis System after 18 h. Image J with the angiogenesis plug-in was used to quantify the angiogenic parameters specified in the figure legends ^57^.

### Sprouting assay in microfluidic device

Microfluidic devices were prepared as previously described ^36^. Briefly, polydimethylsiloxane was cast into previously established 3D printed molds and bonded to glass coverslips with a plasma etcher. Devices were treated with 0.01% (w/v) poly-L-lysine and 0.5% (w/v) L-glutaraldehyde sequentially to promote extracellular matrix attachment to the polydimethylsiloxane housing. Stainless steel acupuncture needles (300 μM, Lhasa OMS) were threaded into each device and type I rat tail collagen (Corning) was prepared as in Doyle ^58^ and polymerized around each needle set for 30 min at 37 °C. Collagen gels were hydrated overnight in EAhy926 media and needles were removed to form 3D hollow channels embedded within the collagen. A 10 μL suspension of cells (2 million/mL) was added to one reservoir of the endothelial channel and inverted for 30 min to allow cell attachment, followed by a second seeding with the device upright. Devices were cultured with continual reciprocating flow utilizing gravity-driven flow on a seesaw rocker plate at 0.33 Hz. Vascular endothelial growth factor (VEGF, Peprotech) was supplemented in culture medium at 10 ng/mL for 48 h, refreshing medium every 24 h. Culture media was further supplemented with EdU for the final 24 h. Tip cell formation was quantified via a custom MATLAB image analysis code with tip cells denoted as any nuclei fully outside of the cell channel edge. Cultures were fixed in 4% paraformaldehyde and permeabilized with a PBS solution containing Triton X-100 (5% v/v), sucrose (10% w/v), and magnesium chloride (0.6% w/v) for 1 h each at room temperature. 4’,6-Diamidino-2-phenylindole (DAPI, 1 μg/mL) was utilized to visualize cell nuclei. For proliferation studies, EdU fluorescent labeling was performed following the manufacturer’s protocol (ClickIT EdU, Life Technologies). DyLight 649 labelled Ulex europaues Agglutinin-1 (UEA, 1:200, Vector Labs) was utilized to visualize endothelial cell location. Fluorescent images were captured on a Zeiss LSM800 confocal microscope.

### Glucose and lactate quantitation

The glucose and lactate quantitation were performed in the culture medium as previously described^23^. Briefly, 100 μL of medium was removed and mixed with 200 μL of 5% HClO_4_, vortexed, and stored at −20 °C until use. On the day of analysis, samples were thawed on ice, vortexed, and centrifuged at 13,000 X g for 5 min at 4 °C. The supernatant was collected and neutralized to pH 7 with saturated K_2_CO_3_ solution. The concentrations of glucose and lactate in the neutralized supernatants were measured using D-GLUCOSE-HK kit (Megazyme) and L-Lactate assay kit (Cayman Chemical), respectively, as per the manufacturers’ protocols.

### Xenograft tumors

SQOR KO mice (both sexes) were given 5 consecutive dosages of 200 μL of tamoxifen (10 mg/mL) in corn oil at an interval of 24 h. On the 8^th^ day, the lower flanks of the mice were implanted with 2 million YUMM 5.2 cells in 200 μL 1X PBS. Mice were sacrificed after 10-12 days and the tumors were excised. Tumor weight was measured and tumor crosssections were prepared for histological and immunohistochemical analyses.

### Immunohistochemistry

Histology were performed at the University of Michigan Centre for Gastrointestinal Research (UMCGR). Tumor tissue and gastrocnemius muscle were fixed in formalin for 24 h, then embedded in paraffin. Sectioning (5 μm thickness) was done using a microtome (Leica RM 2125 RT). The Discovery Ultra (autostainer) platform was used for immunohistochemistry, which involves the following steps. Deparaffinization of tumor and muscle section was performed for 24 min at 69 °C in EZ prep solution. Next, antigen retrieval was done in cell conditioning 1 buffer (CC1) at 95 °C for 40 min. Blocking was done by inhibitor chromoMap for reducing in endogenous peroxidase activity at 37°C for 8 min. Sections were stained with anti-K67 (1:250) and anti-CD31(1:200) antibody for 60 min at 40 °C and washed with reaction buffer (Tris based buffer with preservatives, pH 7.4-7.8). Sections were treated with OmniMap anti-Rb HRP and incubate for 28 min. Next, the sections were treated with ChromoMap DAB solution for 5 min. Sections were counterstained by hand with 5 quick dips in hematoxylin stain, Gill II and any residual stained was removed with the wash buffer. Sections were dehydrated in (95% alcohol, 5mins) before mounting with Permount Mounting Medium. Slides were scanned using a Pannormic scanner and analyzed using case viewer (Caliper Life Sciences).

#### Metabolomics Sample Preparation

For *in vitro* extracellular (media) and intracellular metabolomic profiling, cells were seeded in triplicates in a 6-well plate at 400,000 cells per well in growth media. A parallel plate for protein estimation and sample normalization was also set. After overnight incubation, the culture media was aspirated and replaced with ‘treatment media’. The cells were then cultured for a further 24 h. Thereafter, for extracellular metabolites, 200 μL of media was collected from each well into a 1.5 mL Eppendorf tube and to that 800 μL ice-cold methanol was added. For intracellular metabolites, the remaining media was aspirated, and samples washed 1x with 1mL cold PBS before incubation with 1 mL ice cold 80% methanol on dry ice for 10 min. Thereafter, cell lysates were collected from each well and transferred into separate 1.5 mL Eppendorf tubes. The samples were then centrifuged at 12,000 x *g*. The volume of supernatant for each experimental condition was determined based on the protein concentration of the parallel plate.

#### Targeted Metabolomics

The collected supernatants were dried using a SpeedVac Concentrator, reconstituted with 50% ^v^/v methanol in water, and analyzed by liquid chromatography-coupled mass spectrometry (LC-MS), as described in detail previously^59^.

### Hind limb ischemia through femoral artery ligation

Surgery was performed at the Physiology Phenotyping Core, University of Michigan. Briefly, *Sqrdl^fl/fl^* and *WB^Cre^Sqrdl^fl/fl^* were anasthesized with isoflurane and a circulating heated waterpad was used to maintain the body temperature. An ~1 cm long incision was made from the knee towards the medial left thigh. Subcutaneous fat tissue surrounding the thigh muscle was gently brushed away using sterile saline moistened fine pointed cotton swabs to reveal the underlying femoral artery. Fine forceps and a fine pointed cotton swab was used to gently pierce through the membranous femoral sheath to expose the neurovascular bundle. Silk suture was used to double ligate the femoral artery proximal to the superficial caudal epigastric artery and transected between the two ligatures. The incision was closed using stainless steel wound clips ^60^.

### Laser Doppler Imaging

Laser doppler imaging was performed at physiology phenotyping core, University of Michigan following IACUC guidelines. Briefly, the rodents under anesthesia was monitored by measuring body temperature with a rectal probe, respiration patterns and tail and ear color for cardiovascular function. Mice were anesthetized, one at a time with an isoflurane precision vaporizer anesthetic machine by first placing them in an induction chamber and introducing isoflurane along with approximately 1 liter of oxygen until the animal was recumbent and breathing slowed. The animal was then removed from the induction chamber and placed on a circulating heating pad designed for rodent surgery and temperature regulation and subjected to pre-surgery anesthesia testing by toe pinch. Fur was either shaved using animal clippers and/or completely removed with depilatory cream. Non-invasive imaging was performed with a Perimed Laser Doppler scanning system with a surface probe over a period of 30 min. Since the scanner does not contact the animal, medium is not applied to the surface. Imaging was performed on the ventral surface of the hind limbs on days 1, 3 and 8 post surgery.

