## Supplementary information for "Sulfide oxidation promotes hypoxic angiogenesis and neovascularization"

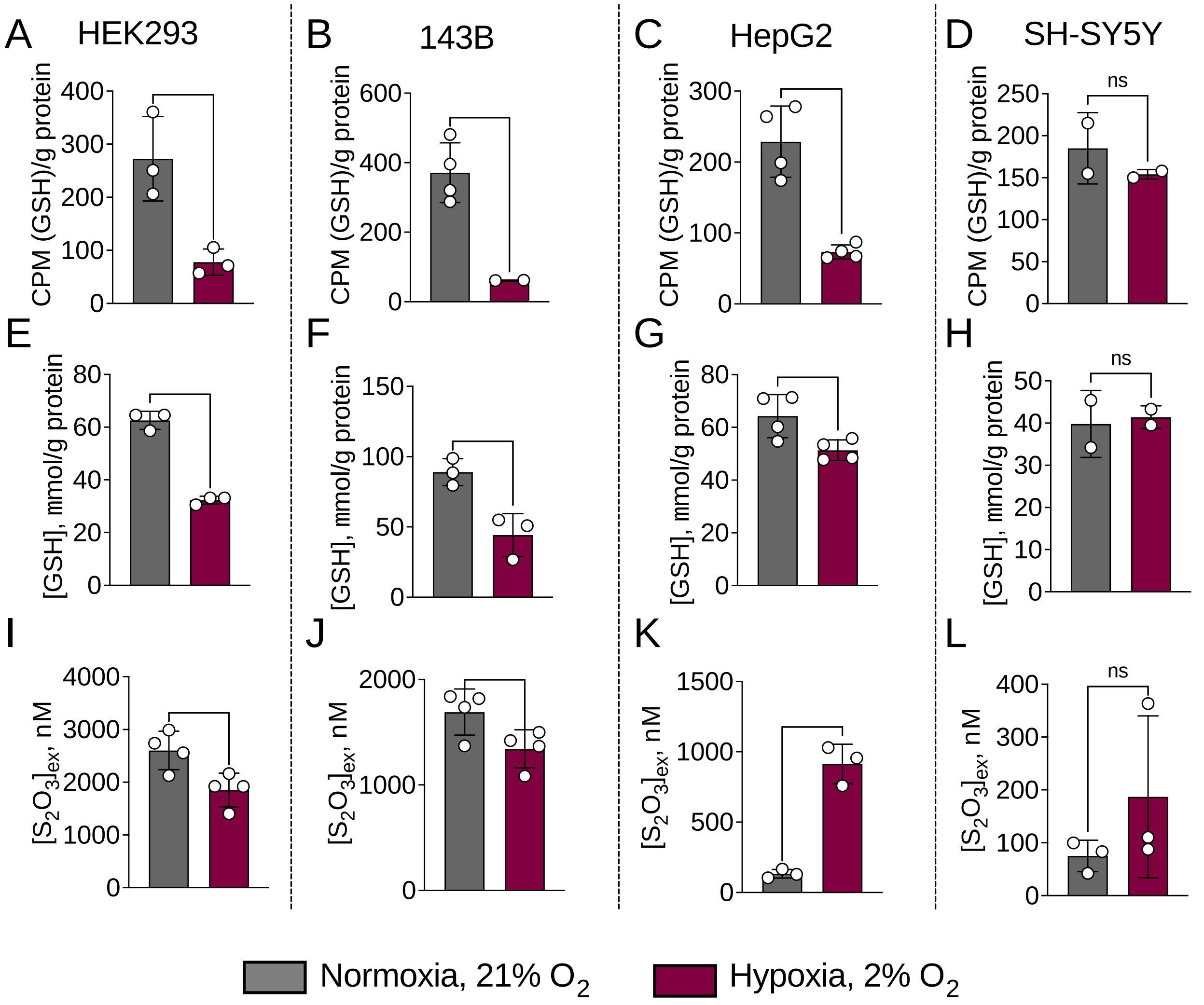


***Supplementary Figure 1. Comparison of transsulfuration pathway activity under normoxia versus hypoxia in non-endothelial cells.*** [$^{35}$S]-methionine incorporation into GSH under 2% (hypoxia) or 21% (normoxia) $O_2$ in: (**A**) HEK293 (**B**) 143B (**C**) HepG2 and (**D**) SH-SY5Y cells. Total GSH content in: (**E**) HEK293, (**F**) 143B, (**G**) HepG2, and (**H**) SH-SY5Y cells. Extracellular thiosulfate $[S_2O_3^{2-}]_{ex}$ accumulation in: (**I**) HEK293, (**J**) 143B, (**K**) HepG2, (**L**) SH-SY5Y. The cells were cultured for either 5 (HepG2) or 24 h (all others). Each data point is from an independent experiment, and the error bars represent ± S.D. *p<0.05 and **p<0.001.

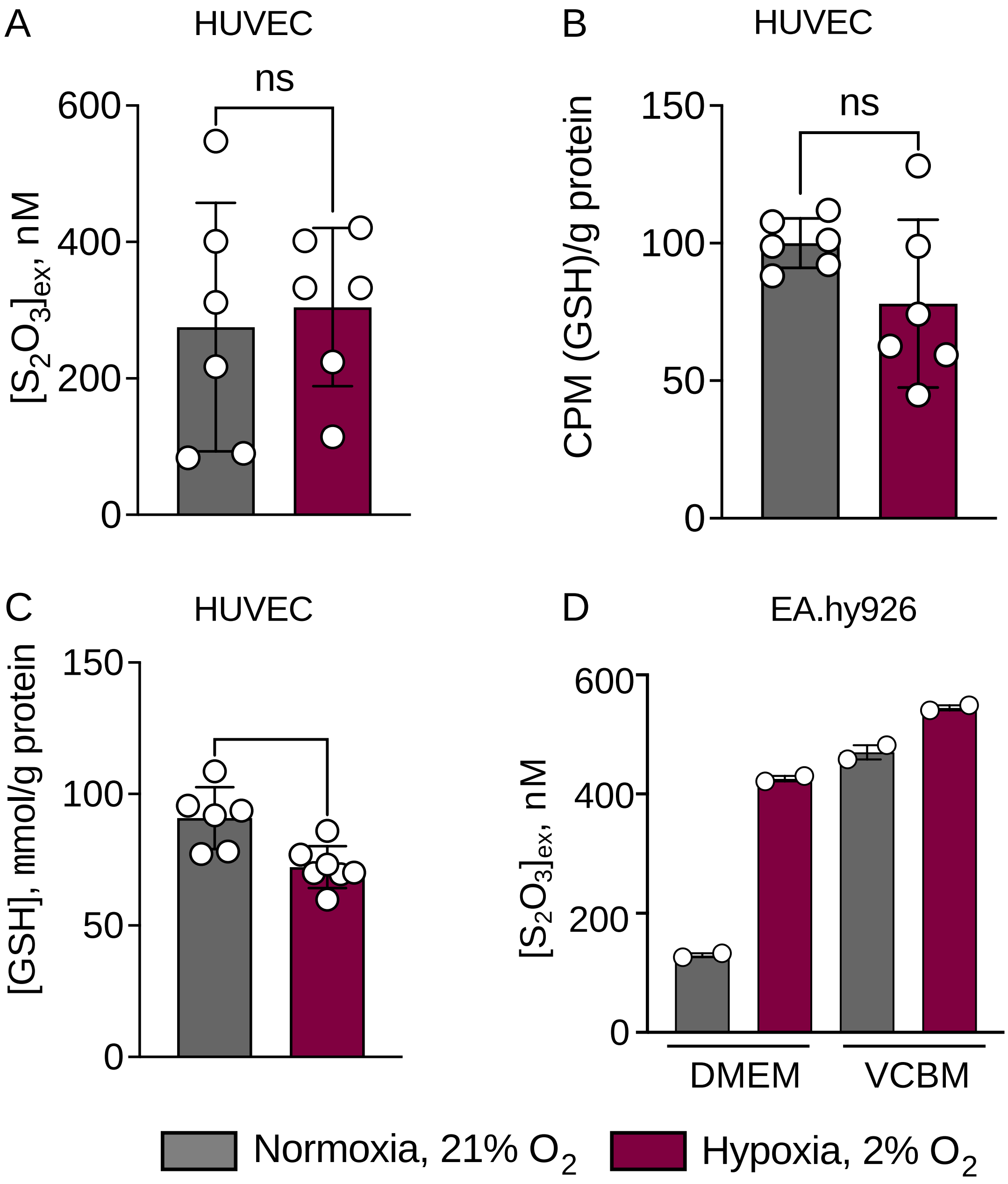


***Supplementary Figure 2. Growth factors influence thiosulfate accumulation.*** (**A**) Thiosulfate levels in conditioned medium, (**B**) [$^{35}$S]-methionine incorporation into GSH, and (**C**) GSH concentration after 24 h of culture of HUVECs in 2% versus 21% $O_2$ in vascular cell basal medium (VCBM) containing a mixture of growth factors (n=3, each in duplicate, **p<0.001). (**D**) EA.hy926 cells cultured in DMEM but not in VCBM showed hypoxic thiosulfate accumulation (n=2). The error bars represent ± S.D.

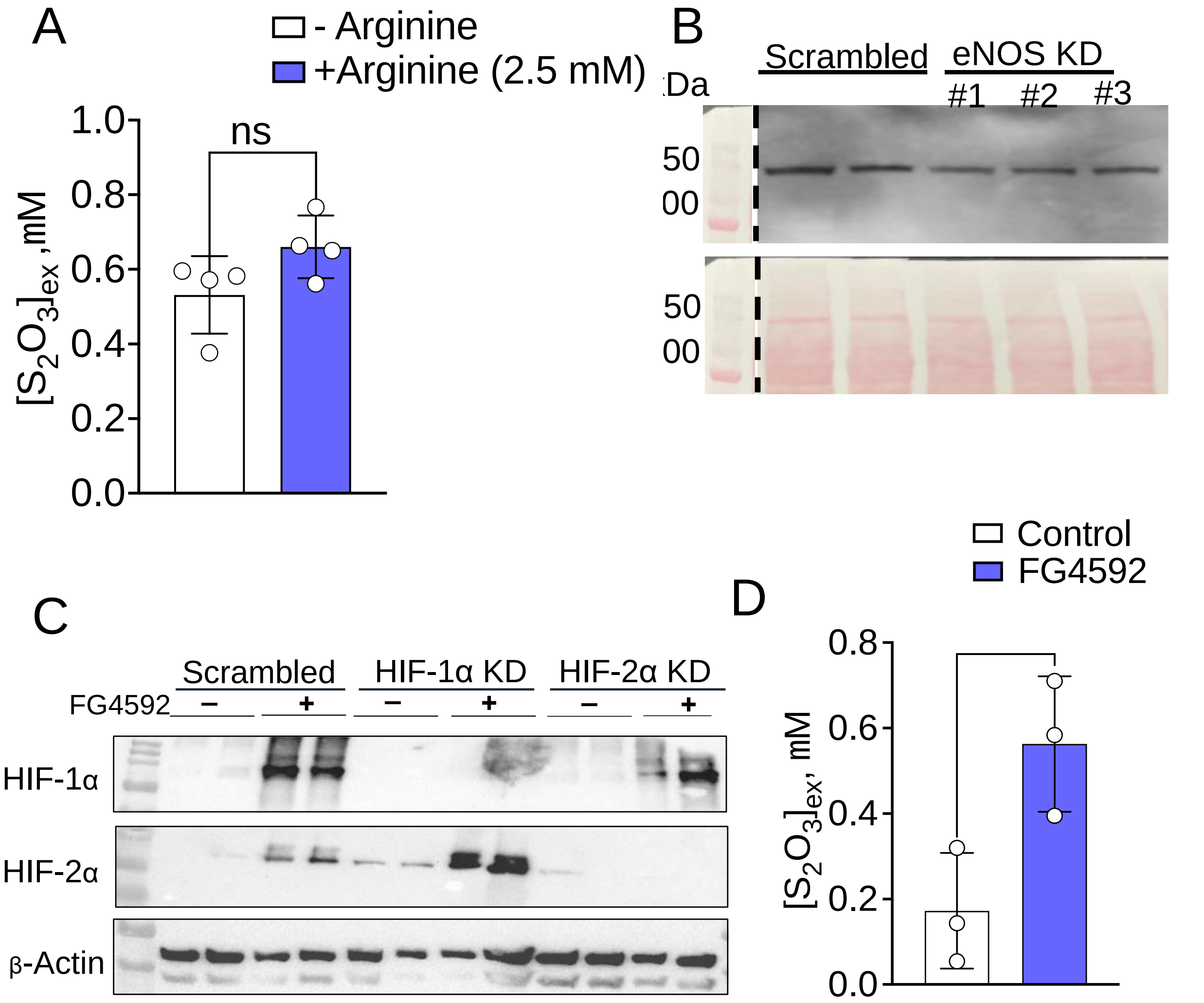


***Supplementary Figure 3. Intersection of •NO and HIF signaling in hypoxic regulation of $H_2S$ homeostasis*** .(**A**) Arginine does not significantly increase extracellular thiosulfate accumulation after 24 h. (**B,C**) Western blot analysis of eNOS (B), HIF1α and H1F2α (C) knockdowns in EA.hy926 cells. Samples were loaded in duplicate. (**D**) FG4592 (roxadustat), a prolyl hydroxylase inhibitor, stabilizes HIF and promotes thiosulfate accumulation under normoxic conditions (24 h). Each data point is from an independent experiment, and the error bars are ± S.D., *p<0.05.

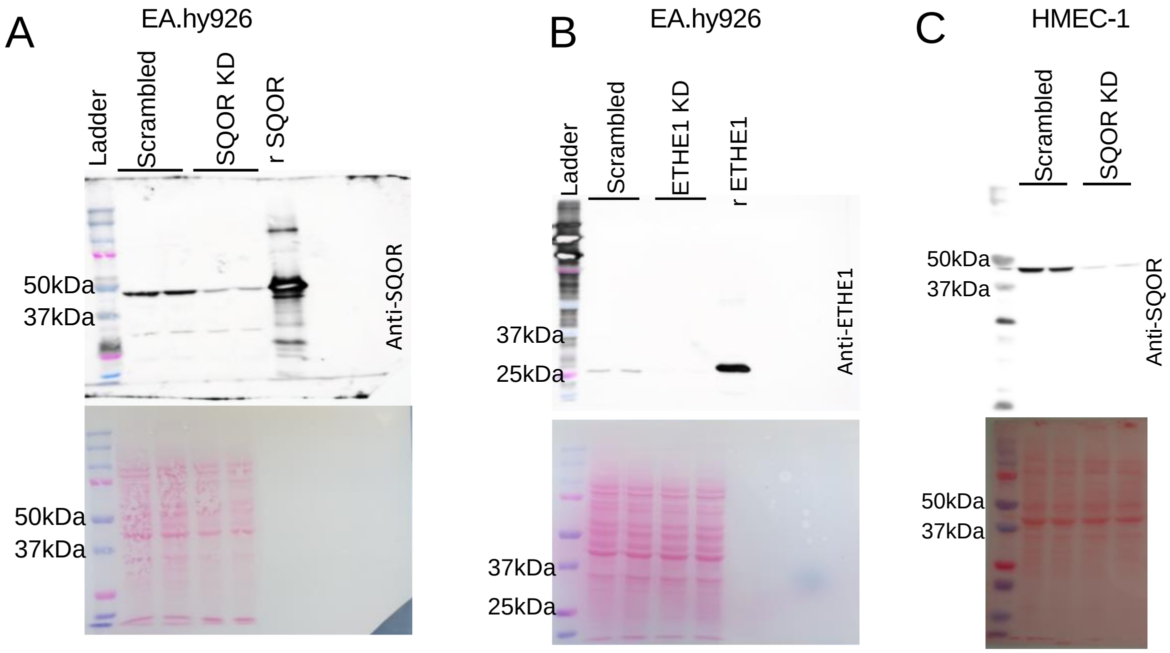


***Supplementary Figure 4. Validation of sulfide oxidation pathway knockdowns.*** **(A,B)** SQOR and ETHE1 knockdown in EA.hy926 cells. (**C**) SQOR knockdown in HMEC-1 cells. rSQOR and rETHE1 denote recombinant human protein standards. Ponceau staining demonstrating equal loading across the lanes (lower panels).

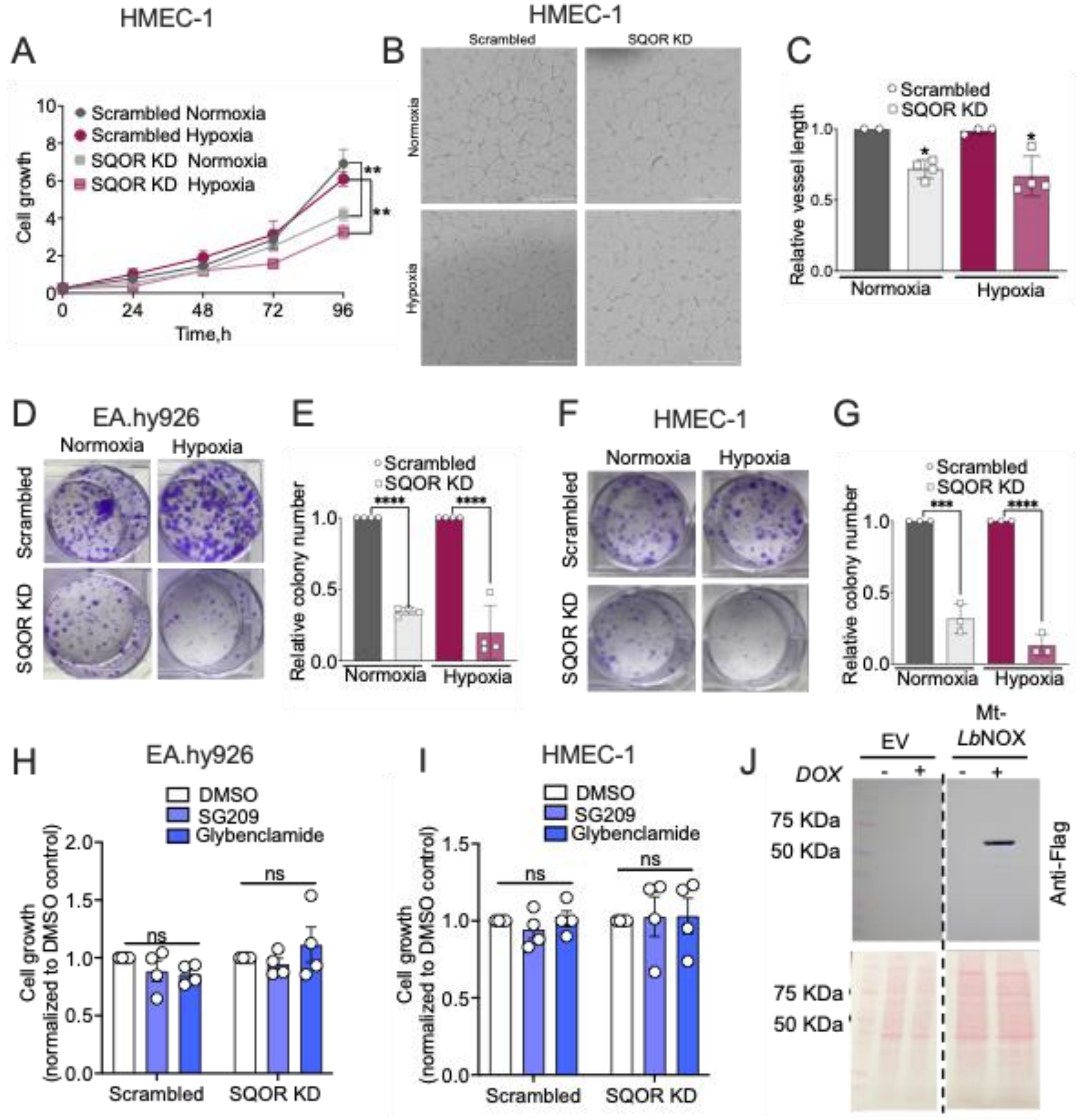


***Supplementary Figure 5. Knockdown of SQOR restricts cell growth.*** SQOR KD lowers HMEC-1 cell proliferation (n=3, **p<0.001.) (**A**), and tube formation (**B, C**) under normoxic and hypoxic conditions (n=3 *p<0.05). (**D,E**) Reduced colony formation (over 10-12 days) was seen in SQOR KD in EA.hy926 cells (D), which was quantitated (n=2 each in duplicate, ****p<0.0001) (E). (**F,G**) Reduced colony formation was seen in SQOR KD in HMEC-1 cells (F), which was quantitated (n=3 , ****p<0.0001) (G). (**H,I**) Effect of the $K_{ATP}$ channel activator SG209 (1 μM), and inhibitor glybenclamide (10 μM) on EA.hy926 (H) and HMEC-1 cell growth (I). (**J**) Expression of mitochondrial *Lb*NOX (Mt-*Lb*NOX) in SQOR KD EA.hy926 cells was validated using an anti-Flag antibody. Error bars represent ± S.D.

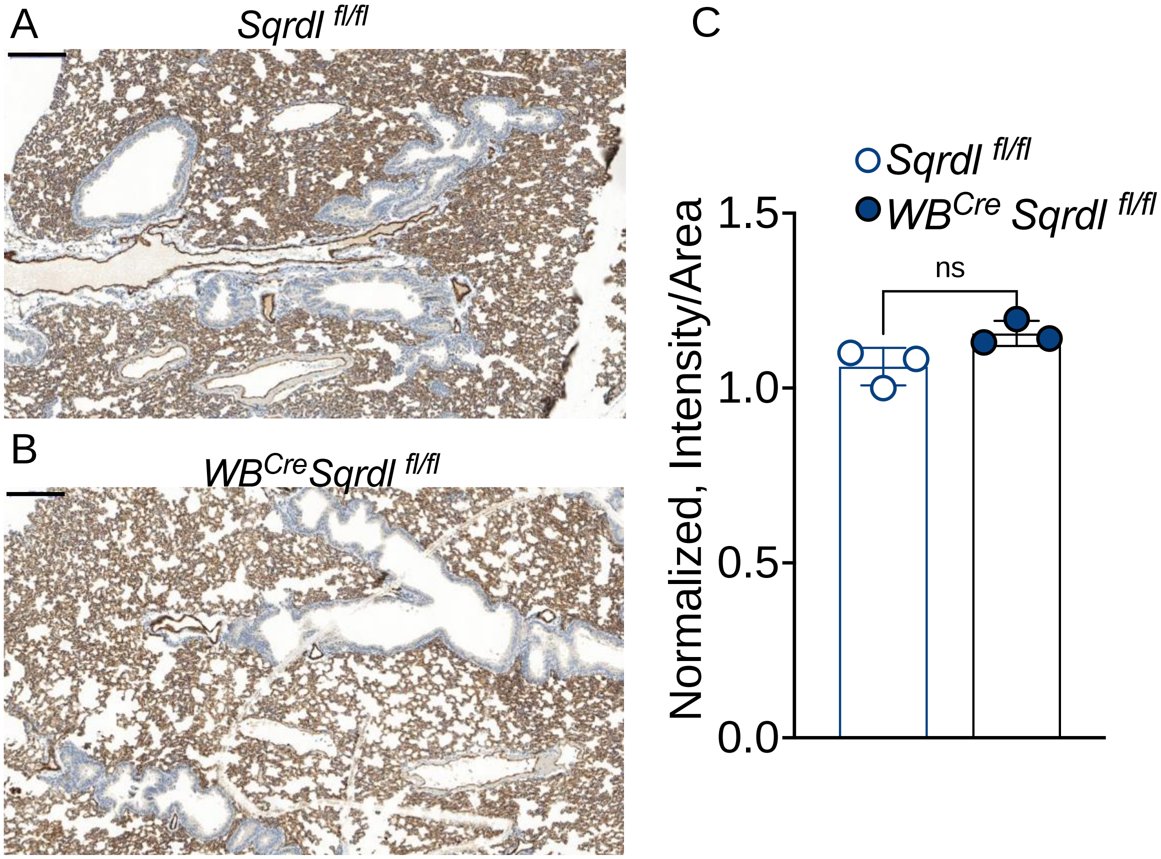


***Supplementary Figure 6. CD31 staining in lung.*** Immunohistochemical analysis of lung cross sections stained with CD31, an endothelial cell marker in control *Sqrdl*$^{fl/fl}$ (**A**) and whole-body SQOR KO mice (WB$^{Cre}$*Sqrdl*$^{lfl/fl}$) (**B**), Quantitation of CD31 staining shows no significant difference (C). Scale bar in A and B represents 200 μm. Error bars in C are ± S.D.

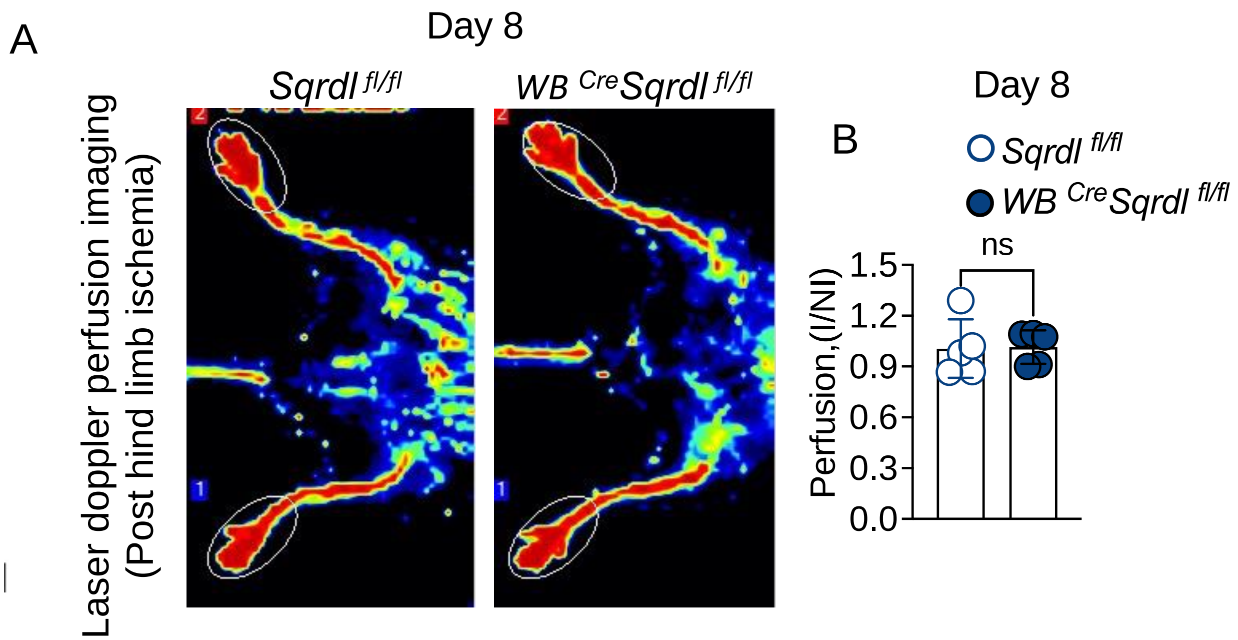


***Supplementary Figure 7.*** **(A)** Laser doppler perfusion imaging on day 8 post hind limb ischemia induction in *Sqrdl*$^{fl/fl}$ and WB$^{Cre}$*Sqrdl*$^{lfl/fl}$ mice. (**B**) No significant differences in blood flow were observed on day 8 following the procedure. Error bars are ± S.D.
